# Wnt-associated adult stem cell marker Lgr6 is required for osteogenesis and fracture healing

**DOI:** 10.1101/2022.12.09.519810

**Authors:** Laura Doherty, Matthew Wan, Anna Peterson, Daniel W. Youngstrom, Justin S. King, Ivo Kalajzic, Kurt D. Hankenson, Archana Sanjay

**Affiliations:** Department of Orthopaedic Surgery, UConn Musculoskeletal Institute, School of Medicine, School of Dental Medicine, UConn Health, Farmington, CT, 06030; School of Dental Medicine, School of Dental Medicine, UConn Health, Farmington, CT, 06030; Department of Reconstructive Sciences, School of Dental Medicine, UConn Health, Farmington, CT, 06030; Department of Orthopaedic Surgery, School of Medicine, University of Michigan, Ann Arbor, MI, 48109

**Keywords:** Fracture healing, bone regeneration, skeletal stem cells, periosteum, Lgr family

## Abstract

Despite the remarkable regenerative capacity of skeletal tissues, nonunion of bone and failure of fractures to heal properly presents a significant clinical concern. Stem and progenitor cells are present in bone and become activated following injury; thus, elucidating mechanisms that promote adult stem cell-mediated healing is important. Wnt-associated adult stem marker Lgr6 is implicated in the regeneration of tissues with well-defined stem cell niches in stem cell-reliant organs. Here, we demonstrate that Lgr6 is dynamically expressed in osteoprogenitors in response to fracture injury. Using an *Lgr6-*null mouse model, we find that *Lgr6* expression is necessary for maintaining bone volume and efficient postnatal bone regeneration in adult mice. Skeletal progenitors isolated from *Lgr6-null* mice have reduced colony-forming potential and reduced osteogenic differentiation capacity due to attenuated cWnt signaling. *Lgr6*-null mice consist of a lower proportion of self-renewing stem cells. In response to fracture injury, *Lgr6-null* mice have deficient proliferation of periosteal progenitors and reduced ALP activity. Further, analysis of bone regeneration phase and remodeling phase of fracture healing in Lgr6-null mice showed impaired endochondral ossification and reduced mineralization. We propose that in contrast to not being required for successful skeletal development Lgr6-positive cells have a direct role in endochondral bone repair.

## Introduction

Insufficient bone healing occurs with high incidence and remains a significant public health concern.^1^ Common comorbidities and risk factors, including advanced age, diabetes, smoking, and inflammatory conditions, exacerbate bone healing abnormalities. In addition, fracture nonunion often requires additional surgical intervention, as there are limited therapeutic options at this stage of repair ^2-4^. The consequence of delayed and failed fracture healing result in an enormous burden to the healthcare system ^1^; therefore, efforts to improve approaches to enhance progenitor-mediated repair are increasingly necessary to advance the fields of orthopedics and regenerative medicine.

Skeletal regeneration leading to scarless tissue formation in adulthood is a complex process that involves the recruitment of progenitor cells from multiple sources. It is orchestrated via various signaling pathway interactions ^5-7^. Following a fracture insult, stem and progenitor cells from the periosteum, bone marrow, and various other sources actively proliferate at the site of injury^8-10^. In this endochondral ossification healing process, these cells differentiate into chondrocytes and osteoblasts to produce a cartilaginous matrix and osteoid, respectively; the generated fracture callus later becomes mineralized and is remodeled to its original architecture. The periosteal layer harbors essential osteochondral progenitors that are robustly activated following the fracture, contributing to the callus formation via endochondral ossification at the central injury area and intramembranous ossification flanking the fracture site ^5,8-11^. Removal of the periosteum and the osteochondral progenitors within severely impair efficient fracture healing ^5,8,10-12^.

Leucine-rich repeat-containing G protein-coupled receptor 6 (Lgr6) is a protein in the G-protein-coupled 7-transmembrane superfamily ^13^. Members of the Lgr family (Lgr4/5/6) are primarily characterized as adult stem cell markers that play a critical role in defining stem and progenitor cell behavior in established stem cell niches ^14-16^. In adult stem cell-reliant organs (e.g., intestine, hair follicle, skin), different Lgrs mark distinct stem cell populations regarding location and function ^14,15^. In some Wnt-driven stem cell compartments, Lgrs can serve an additional function as receptors in the Wnt/β-catenin pathway ^17,18^; in this context, binding of soluble R-spondins (Rspo1-4) to Lgrs prolongs the presence of Frizzled (Fzd) receptors on the cell surface, as Lgrs interact with components of the Wnt signalosome. Recent work has also indicated that Lgr6 is dynamically expressed in mouse and human-derived mesenchymal stem cells and plays a role in osteogenesis ^19-23^. However, very little is known about its role in bone regeneration, prompting our current studies on this gene in osteoblast differentiation and bone healing.

Here, we report that Lgr6 is dynamically expressed in osteoprogenitors in response to fracture injury. We found that ablation of *Lgr6* reduces the bone volume in mature mice of either sex. *Lgr6* is required for osteogenic differentiation *in vitro*, and its expression confers colony-forming potential in isolated mesenchymal progenitors from adult mice. Additionally, endochondral fracture healing and callus mineralization are significantly impaired in *Lgr6-*null mice. We propose that *Lgr6*-positive cells have a direct role in endochondral bone repair and that *Lgr6* functions during postnatal bone regeneration, in contrast to not being required for successful skeletal development.

## Results

### Increased Lgr6 expression in a mouse model with enhanced periosteal response to injury

Previous studies from our group have used a knock-in mouse model with a global increase in PI3K activity (gCbl^YF^) to evaluate various mechanisms of bone healing and skeletal cell differentiation ^24-29^; due to a point mutation, these mice lack the interaction between the adaptor protein Cbl and the p85 regulatory subunit of PI3K. We reported an expansion of periosteal cells flanking the fracture site in the early response to bone injury using gCbl^YF^ mice ^30^, where they exhibit increased periosteal activation post-fracture, increased bony callus formation, and increased periosteal osteogenic potential *in vitro*. The three-fold increase in periosteal thickness during mesenchymal progenitor activation correlates with a subsequent increase in osterix- and ALP-positive bone lining cells ^30^. Bulk mRNA-sequencing was employed to identify novel transcripts upregulated in gCbl^YF^ periosteal cells compared to controls that would account for the robust periosteal expansion. Our analysis found that many cWnt/β-catenin pathway-related genes were upregulated in gCbl^YF^-derived periosteal cells compared to controls (Fig.1a). These genes included Wnt ligands (*Wnt10a, Wnt10b, Wnt5a*), Wnt-associated receptors (*Frzb, Lrp6*), Wnt inhibitors/antagonists (*Sfrp2*, and *Notum*), as well as Wnt target gene (*Sp7*).

One of the most significantly upregulated genes in this data set was *Lgr6* (5.82-fold change, P-adj =1.75 E-156). The Lgr6 protein is an adult stem cell marker ^14-16^ and atypical G-protein-coupled receptor (GPCR) ^13^ that modulates the activity of the canonical Wnt (cWnt) pathway ^18,31^. Among the other members of this receptor family, *Lgr5* expression was downregulated in gCbl^YF^-derived cells compared to controls, and there was no significant change in the expression of *Lgr4*. The differential expression of these Lgr family members in the gCbl^YF^ periosteal cells was confirmed by qRT-PCR (Fig.1b).

**Figure 1.**
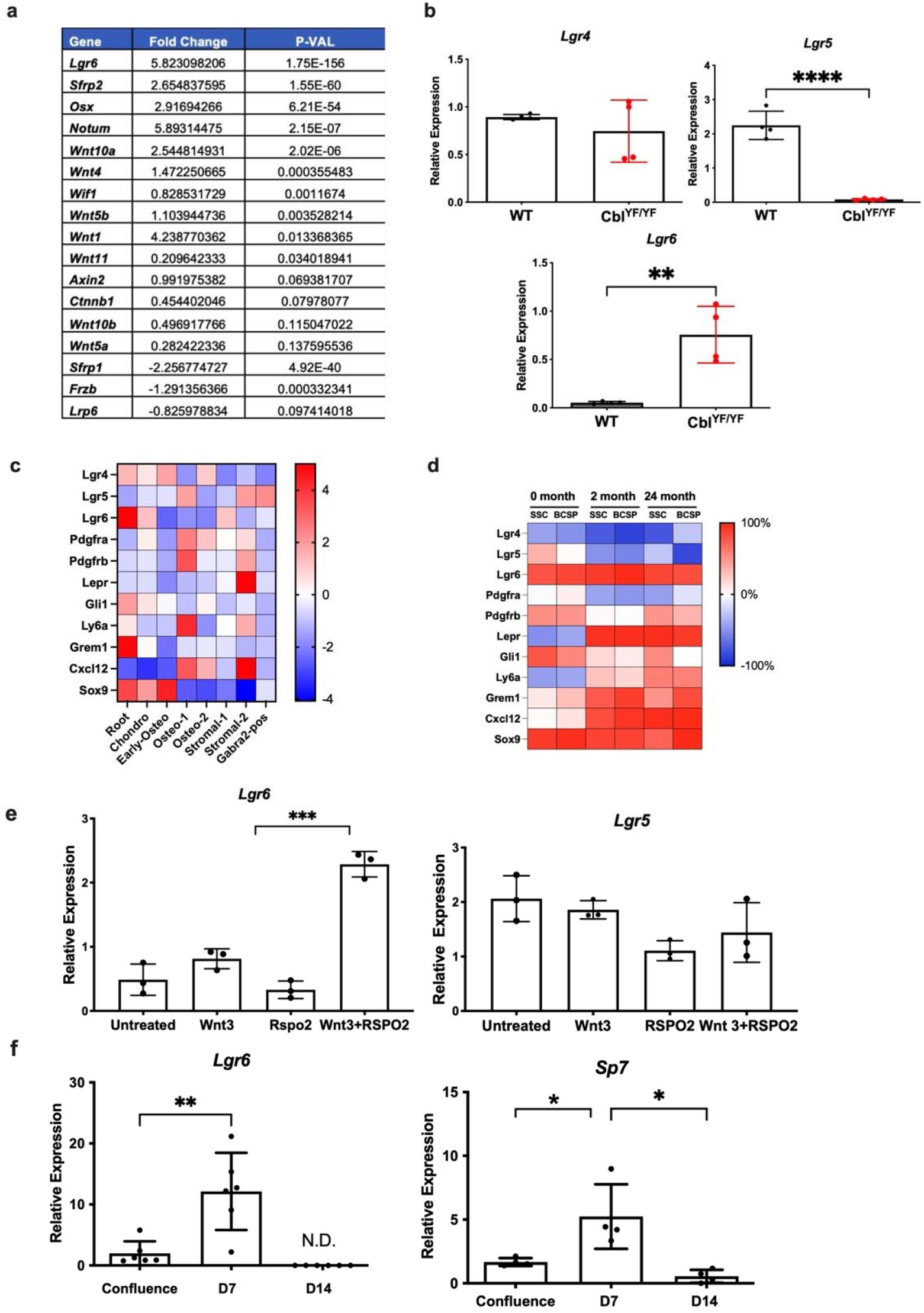
**a**. Wnt pathway-related genes identified to be differentially expressed in gCbl^YF^-derived periosteal cells compared to controls. **b**. To determine Lgr6 expression specifically in self-renewing stem cells (SSCs) in bone marrow, we analyzed a recently published Smart-seq2 single RNA Seq data from long bones derived from postnatal day 3, 2-month, and 24-month-old mice^33^. Relative expression analysis in the SSC population showed *Lgr6* was among the highly expressed known stem and progenitor cell markers in osteochondrogenic lineage. **c**. From the same study, we also analyzed select genes from microarray data from a skeletal stem cell (SSC; CD51^+^ THY1^-^6C3^-^CD105^-^) and bone-cartilage-stromal progenitor (BCSP; CD51^+^THY1^-^ 6C3^-^CD105^+^) population derived from 0-, 2- and 24-month-old mice^33^. Our analysis showed that compared to some of the other SSC markers, Lgr6 expression was minimally changed with aging. **d**. Periosteal cultures were grown to confluence, and the expression of Lgr family members was analyzed by qRTPCR. **e**. Periosteal cultures were grown to confluence and treated with Wnt3a (50 ng/mL) or a combination of Wnt3a and Rspo2 (50 ng/mL). qRTPCR analysis was performed to detect the expression of *Lgr6 and Lgr5*. **f**. Periosteal cultures were grown to confluence. qRTPCR analysis shows *Lgr6* and *Sp7* expression in isolated periosteal cells is significantly reduced upon 7 days of osteogenic differentiation. n=3 *p<0.05. A representative of 2 experiments is shown.

Lgr6 marks distinct stem cell populations in terms of location and function in the hair follicle, nail bed, and lung ^16,21,32^. To determine Lgr6 expression specifically in self-renewing skeletal stem cells (SSCs), we analyzed a recently published single RNA Seq data from the long bones derived from postnatal day 3 (0-month), 2-month, and 24-month-old mice ^33^. Relative expression analysis in the SSC population showed that *Lgr6* was highly expressed (log fold change =1.447, P-adj = 0.000426) in a cluster that also expressed other known stem and progenitor cell markers in the osteochondrogenic lineage, including *Sox9* (log fold change = 4.688, P-adj = 0.0428) *Grem1* (log fold change = 2.808, P-adj =0.000831) and *Gli1* (log fold change = 0.880, P-adj = 0.749) (Fig. 1c). We also analyzed the microarray data of SSCs and bone–cartilage– stromal progenitors (BCSPs) derived from young, mature, and aged mice ^33^. Among these populations, *Lgr6* expression was higher compared to Lgr4 and Lgr5. Furthermore, *Lgr6* expression minimally changed with age compared to other SSC markers (Fig. 1d). Together, this data indicates that *Lgr6* is expressed in mesenchymal stem/progenitor cells residing in long bones.

### *Lgr6* expression correlates with osteogenic differentiation of osteochondral progenitor cells

Wnt signaling regulates osteogenic differentiation ^34^, and Lgrs are known targets of Wnt signaling ^17,35^. We found that the combined treatment of Wnt3a and Rspo2 of wild-type periosteal cells increased *Lgr6* expression 2.5-fold more than treating cells with Wnt3a or Rspo2 alone; in contrast, no upregulation was seen for *Lgr5* expression (Fig. 1c). *Lgr6* is dynamically expressed in calvaria- and bone marrow-derived mesenchymal cells during osteogenic differentiation *in vitro* ^19,20^. We examined *Lgr6* expression in primary periosteal cells since the periosteum harbors osteochondral progenitors and is essential to homeostatic bone formation and fracture healing ^5,8,9^. In the early phase of osteogenic differentiation, we observed an increase in *Lgr6* expression with a corresponding increase in *Sp7* expression; Levels of *Sp7* and *Lgr6* expression are downregulated after 14 days of osteogenesis (Fig. 1e). This expression pattern mirrors studies showing that *Lgr6* is upregulated in mesenchymal cells upon osteoinduction but decreases during osteoblastic differentiation and maturation ^19,20^. *Lgr4*, another LGR family member studied in bone-specific contexts, does not demonstrate this dynamic expression ^31,36^. Together, these results suggest that Lgr6 has a specific temporal role in osteoblast differentiation.

### *Lgr6* expression is increased in response to fracture healing

We determined Lgr6 expression in a femur fracture model in the context of bone injury. For this purpose, mice with an inactivating EGFP-ires-CreET2 insertion into endogenous *Lgr6* allele were used ^16,19^. *Lgr6*^*EGFP-ires-CreERT2/+*^ mice were bred with a *tdTomato* fluorescent reporter mouse (*Lgr6*^*EGFP-ires-CreERT2/+*^; *Ai9*). Using a short chase following a tamoxifen injection, we found that in uninjured bones, scarce tdTomato+ cells were visible by microscopy only in the bone marrow compartment, and no tdTomato+ cells were found on the periosteum (Fig. 2a and data not shown). Five days after the injury, the periosteum was thickened near the fracture site, and there was an increase in the number of tdTomato+ cells in the cambium layer of the periosteum. Fourteen days post fracture, we found td Tomato+ cells in the fracture callus. However, no tdTomato+ chondrocytes within the callus were visible. Immunostaining with anti-osterix antibodies showed the colocalization of tdTomato+ cells with osterix+ cells in the callus (Fig. 2g-i). Together, these results indicated that in response to bone injury, within the fracture callus, the number of Lgr6+ progenitor cells and their progeny was increased on newly formed bone; many of the tdTomato+ cells were also osterix+.

**Figure 2.**
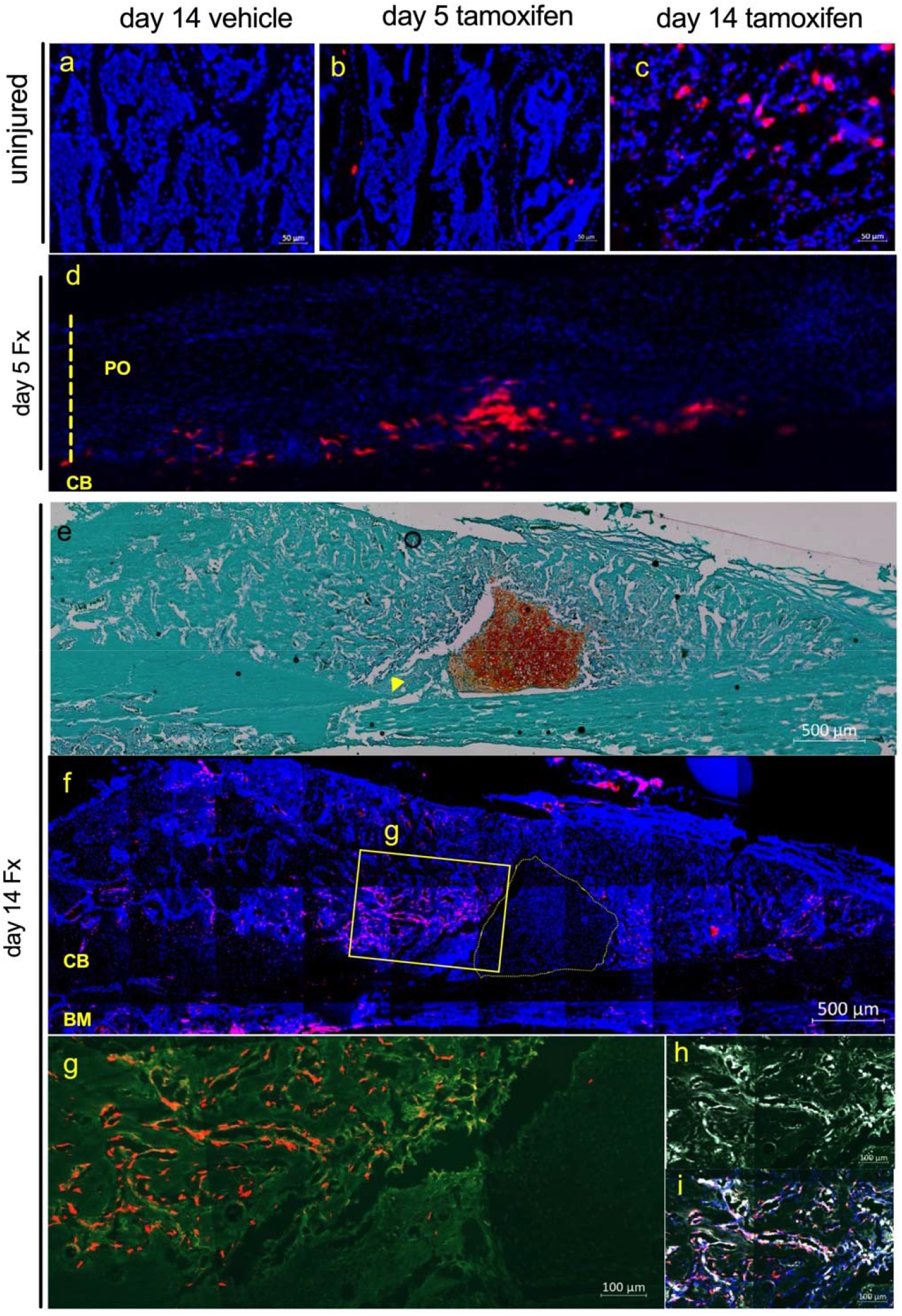
Lgr6 expression in injured bone. *Lgr6*^*EGFP-ires-CreERT2*^*/*^*+*^; Ai9/^+^ mice were injected with (**a**) vehicle or tamoxifen (b-h) and sacrificed for histological assessment 5- and 14-days after injection. **b** and **c**. Progeny of Lgr6+ progenitors are labeled tdTomato+ within the non-injured bone marrow. **d**. Five days post-fracture presence of the progeny of Lgr6+ progenitors labeled tdTomato+ within select areas of the thickened cambium layer. The dotted line shows thickened periosteum. **e**. Safranin O and Fast green staining of sections. Arrow indicates fracture site. 14-day post fracture. **f**. Progeny of Lgr6+ progenitors labeled tdTomato+ within select fracture callus 14-day post fracture. The dotted line indicates chondrocytes. **g**. Magnified boxed area. The bone structure is indicated by green autofluorescence. **h**. Osterix+ cells. **i**. Colocalization of tdTomato+ and osterix+ cells within the fracture callus. PO=periosteum, CB=cortical bone., BM=bone marrow. 5 days post-fracture. n= 5 mice/condition. Representative images are shown.

### Lgr6-null mice are osteopenic

We utilized previously described *Lgr6-*null mice to determine a functional role for *Lgr6* in bone ^16^. We confirmed that *Lgr6*-null mice lack transcriptional activity of the *Lgr6* gene in long bones, and Lgr6 protein was not detected in lysates of bone marrow stromal cell cultures derived from *Lgr6*-null mice (Supplementary Figure 1a and b). While Lgr4-null and Lgr5-null mice have severe skeletal abnormalities and neonatal lethality ^37,38^, our results show that *Lgr6-*null mice develop into healthy, fertile adults with no changes in body weight or mass at 5 months of age (Supplementary Figure 1c), suggesting that Lgr6 is not as developmentally important as Lgr4 or Lgr5. Lgr6-null mice do not have significant differences in bone volume at 3-months; however, at 5-months, both male and female mice show decreased bone volume (data not shown). Microcomputed tomography (μCT) was used to evaluate trabecular and cortical bone parameters of the right distal femur from mice at all time points for both sexes. At five months of age, *Lgr6*-null mice had a decrease in trabecular BV/TV compared to their control sex-matched littermates (BV/TV, males = −49%, P=0.035; females = −85% P=0.045) (Fig.3a). The decreased

BV/TV corresponded to decreased trabecular number (45% and 35% decrease in male and female mice respectively) and increased trabecular spacing (males= 2-fold; females=1.6-fold); trabecular thickness was not changed in males, and there was a 20% increase in *Lgr6*-null females (P= 0.0376) (Fig. 3b; Table1). Compared to controls, in *Lgr6*-null mice, the cortical porosity was decreased (Ct. Po, males 34.4%, P=0.0001; females 27%, P= 0.0087), while no significant differences were identified in the cortical bone area or thickness (data not shown). Despite a decrease in the trabecular bone volume, there were no changes in *Col1a, Runx2*, or *Axin2* gene expression in the femurs of 5-month-old *Lgr6-*null mice (Supplementary Figure 1d).

**Figure 3.**
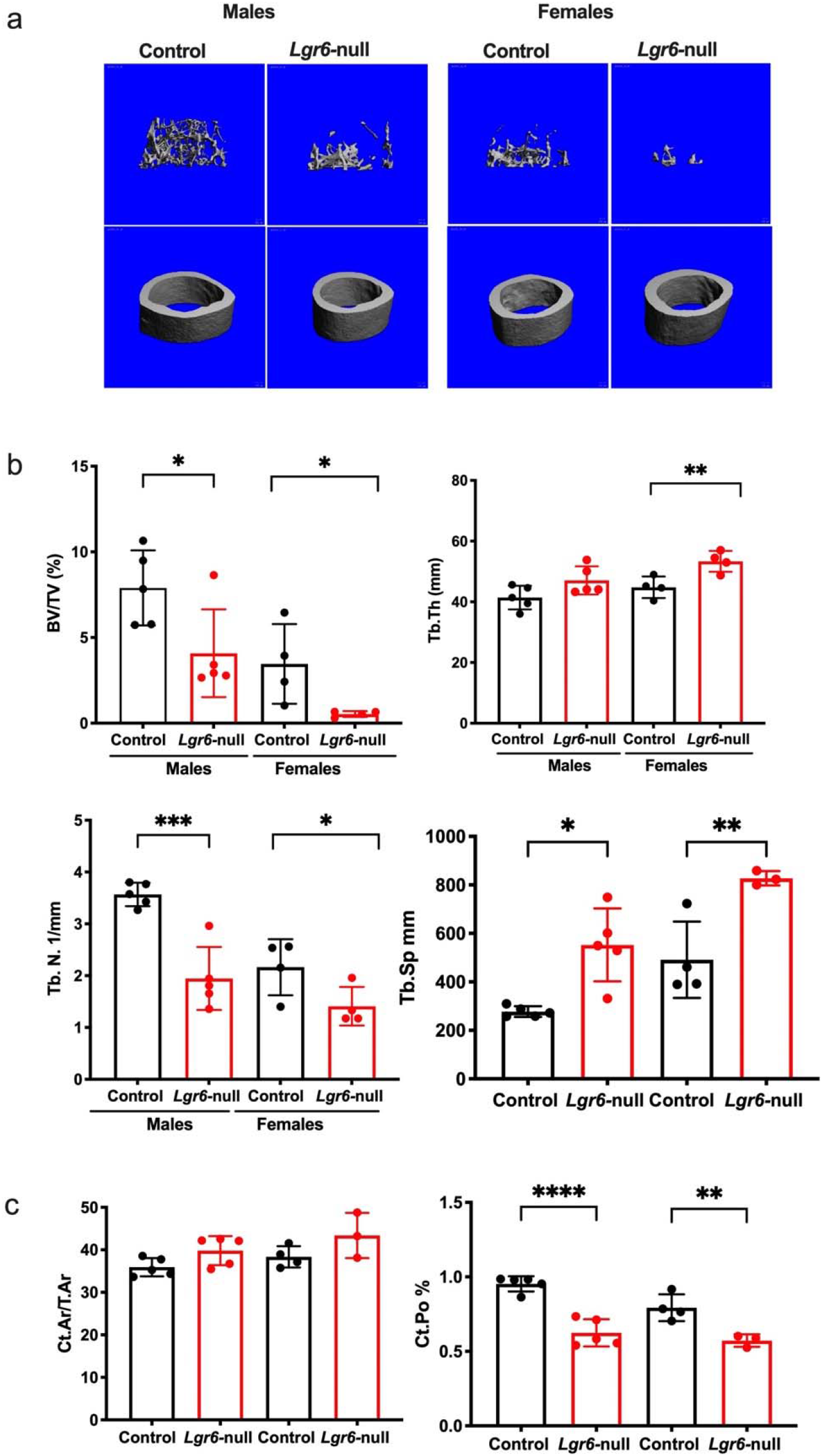
*Lgr6-*null mice are osteopenic. **a**. Representative 3D reconstructed μCT images of femoral trabecular bone from 3-and 5-month-old mice from each sex and genotype **b-g**. Quantified trabecular and cortical μCT parameters between genotypes: **b**. bone volume/tissue volume (BV/TV), **c**. trabecular number (Tb.N.), **d**. trabecular separation (Tb. Sp), **e**. trabecular thickness (Tb.Th.), **f**. Cortical area/tissue area (Ct. Ar./T.Ar.) **g**. Cortical porosity (Ct. Po). n= Controls: 5 males and 4 females; *Lgr6*-null: 5 males and 4 females. Quantified bone parameters were analyzed by Student’s t-test; sexes were analyzed separately. *p<0.05, **p <0.001, ***p <0.0001 Scale bar 1mm.

**Table 1.** Femoral microarchitecture assessed by μCT of 5-month-old male and female mice. μCT was performed on distal femurs for trabecular bone and the femoral midshaft for cortical bone. Values are means ± SD. *Significantly different compared to controls by one-way ANOVA with Tukey’s post-hoc test. *\*p* < 0.05, \**\*p* < 0.01, \**\*\*p* < 0.001,

|  | Male |  | Female |  |
| --- | --- | --- | --- | --- |
|  | Control | *Lgr6*-null | Control | *Lgr6*-null |
|  | Femoral Trabecular Bone |  |  |  |
| BV/TV (%) | 7.90 ± 2.19 | 4.08 ± 2.56 * | 3.46 ± 2.33 | 0.535 ± 0.167* |
| BV (mm<sup>3</sup>) | 0.208 ± 0.084 | 0.105 ± 0.080 | 0.080 ± 0.067 | 0.0132 ± 0.005 |
| TV (mm<sup>3</sup>) | 2.56 ± 0.381 | 2.45 ± 0.243 | 2.41 ± 0.154 | 2.46 ± 0.110 |
| BS (mm) | 12.8 ± 3.98 | 5.63 ± 4.04* | 4.67 ± 3.55 | 0.606 ± 0.242* |
| BS/BV | 65.2 ± 7.04 | 57.1 ± 3.39 | 64.8 ± 9.42 | 48.2 ± 6.59 |
| Trab. Spacing | 277 ± 22.0 | 552 ± 150* | 491 ± 158 | 827 ± 29.8** |
| Trab. Number | 3.57 ± 0.225*** | 1.95 ± 0.608 | 2.16 ± 0.541 | 1.41 ± 0.373* |
| Trab. Thickness | 41.4 ± 3.89 | 47.1 ± 4.65 | 44.8 ± 3.52 | 53.3 ± 3.44** |
|  | Femoral Cortical Bone |  |  |  |
| Tt. Ar (mm<sup>2</sup>) | 1.67 ± 0.149 | 1.37 ± 0.163 | 1.50 ± 0.111 | 1.47 ± 0.142 |
| Ct. Ar (mm<sup>2</sup>) | 0.144 ± 0.007 | 0.147 ± 0.0123 | 0.151 ± 0.007 | 0.170 ± 0.020 |
| Ct. Ar/Tt. Ar | 35.9 ± 2.15 | 39.8 ± 3.43 | 38.4 ± 2.51 | 43.4 ± 5.31 |
| Ct. Th(mm) | 0.144 ± 0.008 | 0.147 ± 0.012 | 0.151 ± 0.007 | 0.170 ± 0.020 |
| Ps. Pm (mm) | 4.76 ± 0.255 | 4.51 ± 0.175 | 4.53 ± 0.136 | 4.40 ± 0.212 |
| Ec. Pm (mm) | 3.85 ± 0.241 | 3.32 ± 0.271 | 3.51 ± 0.170 | 3.33 ± 0.289 |
| Porosity (%) | 0.953 ± 0.051 | 0.625 ± 0.091*** | 0.793 ± 0.090 | 0.572 ± 0.041* |

### Skeletal progenitors isolated from *Lgr6-null* mice have reduced colony-forming potential and reduced osteogenic differentiation capacity due to attenuated cWnt signaling

Using *ex vivo* cultures, we evaluated the colony-forming capabilities of mesenchymal cells harvested from bone marrow and periosteum to assess the cellular basis for the decreased bone volume. Ex vivo, periosteal progenitors derived from *Lgr6-*null mice also showed a significant decrease in CFU-F (Control = 56.3, *Lgr6*-null = 26.6, P=0.001) and CFU-AP+ (Control= 31.6, *Lgr6*-null =12.6, P= 0.0005) (Fig 4a). We also found that at 3-and 5 months of age, there was a 40% and 36 % decrease in CFU-Fs from *Lgr6-*null bone marrow-derived cells, indicating reduced *in vitro* progenitor potential. In addition, *Lgr6-*null cells also had a significant deficiency in CFU-ALP, which was more pronounced with age. While cells from control animals successfully formed numerous ALP+ colonies, in *Lgr6*-null cultures, there was a 50% decrease in CFU-AP at 3 months. Remarkably, *Lgr6-*null cultures derived from 5-month-old mice had only 0-2 CFU-ALP+ colonies (Fig. 4a).

**Figure 4.**
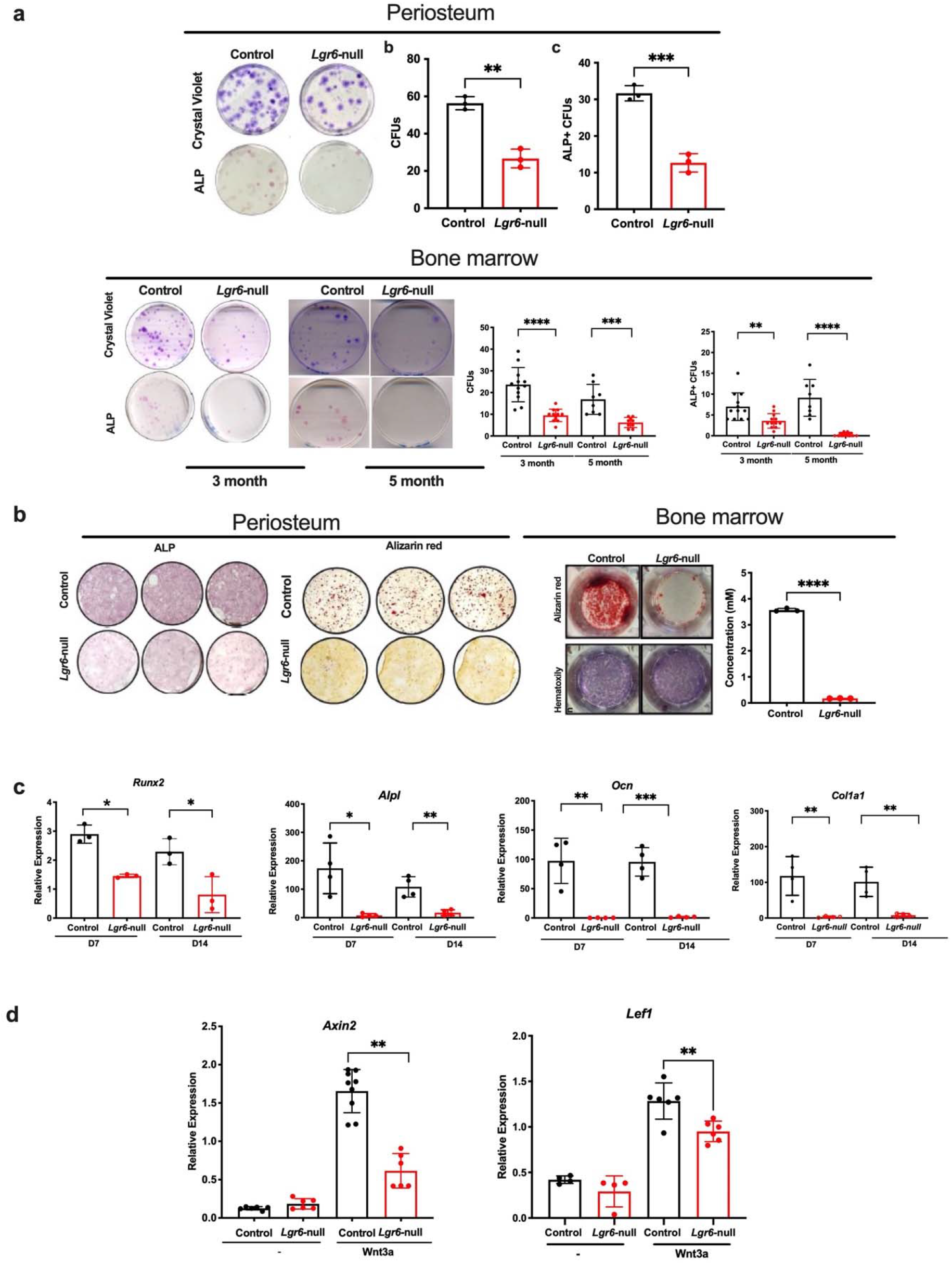
Mesenchymal progenitors isolated from *Lgr6-*null mice have reduced colony-forming potential, reduced osteogenic differentiation capacity, and attenuated response to cWnt signaling. **a**. Periosteum and Bone marrow-derived cells from *Lgr6*-null and control mice were plated for CFU assays and stained with crystal violet staining to identify colony formation; colonies were stained alkaline phosphatase to assess osteogenic potential. Colonies were quantified. Mice of both sexes were used. For periosteal cultures are derived from 3-month-old males and females. Cells for each sample were pooled from two individual mice of the same sex. For BMSCs Control, n=8-12 (4-6 males; 4-6 females) *Lgr6*-null, n=9-12 (4-6 males; 5-6 females). **c**. Periosteal or BMSCs from *Lgr6-*null and control mice underwent *in vitro* osteogenesis for 21 days. Periosteal cells were stained for ALP and with alizarin red. BMSCs were stained for alizarin red-S (upper panel), followed by hematoxylin (lower panel). Quantitation of alizarin red-S dye. **d**. qRT-PCR analyses of osteogenic markers at day 7 and day 14 of osteogenesis of BMSCs. The osteogenic differentiation experiment was performed in triplicates and repeated 3 times. A representative of 3 experiments is shown. **e**. BMSCs grown to confluence were serum starved for 24h and then treated with cWnt3a (100ng/ml) for 6 h. Expression of *Axin2* and *Lef1* was examined. A representative of 2 experiments is shown. Data were analyzed by Students’ t-test; *p<0.05, **p <0.01; ***p<0.0001.

Strikingly, *Lgr6*-null-derived periosteal cells also show severely impaired osteogenic differentiation as evaluated by ALP staining and alizarin red staining of calcium deposits (Fig. 4b). *Lgr6-*null-bone marrow stromal cells from 3-month-old mice also exhibit significant deficiencies in mineralization, while cells from control animals of the same age successfully differentiate into mature, mineral-depositing osteoblasts after 21 days of differentiation (Fig. 4b). This impaired osteogenic capacity is also evident with decreased expression of osteogenic markers, including *Runx2, Alpl, Ocn*, and *Col1a* (Fig. 4c and Supplementary Figure 2). Consistent with the role of Lgr6 in modulating Wnt signaling, we found that compared to controls, *Lgr6*-null cultures exposed to Wnt3a had 67% and 25% decrease in *Axin2* and *Lef1* expression, respectively (Fig. 4d). We confirmed no changes in the potential to differentiate toward the chondrocytic lineage (Supplementary Figure 2). As reduced bone mass could be due to increased osteoclast activity, we assessed *in vitro* osteoclastogenesis assays using bone marrow-derived monocytes (BMMs). We found no differences in the number of osteoclasts in cultures from *Lgr6-*null mice and controls (Supplementary Figure 2).

### *Lgr6*-null mice consist of a lower proportion of self-renewing stem cells

We used the flow cytometry technique to discern further how *Lgr6* modulates progenitor cell populations. We utilized cell surface antigen combinations and gating strategies developed in the Chan and Longaker laboratories to identify skeletal stem cells and their downstream bone, cartilage, and stromal progenitors ^39^. Using this strategy, we evaluated the presence of three distinct populations of skeletal progenitors, which exist in a differentiation hierarchy: self-renewing stem cells (SSC, CD45−Ter119−CD31 −Thy1−6C3−CD51+CD105−CD200+) at the apex and two populations of downstream, more committed progenitors pre bone–cartilage– stromal progenitor (pre-BSCP, CD45−Ter119−CD31 −Thy1−6C3−CD51+CD105−CD200-) and bone–cartilage–stromal progenitors (BSCP, CD45−Ter119−CD31−Thy1−6C3−CD51+) (Fig. 5). Our analysis showed that bone marrow and periosteal cells isolated from *Lgr6-*null mice display a significant shift in the proportion of these populations compared to controls; specifically, a smaller percentage of the subpopulation of self-renewing SSCs in bone marrow (SSC, Control 0.172% SD± 0.38, *Lgr6*-null 0.073 SD ± 0.019, P = 0.0072) and periosteum (SSC, Control 1.805% SD± 0.058, *Lgr6*-null 1.046 SD ± 0.461, P= 0.019). Surprisingly, we found an increase in the percentage of Pre-BSCP in *Lgr6*-null samples (Pre-BSCP, Control =16.8%, *Lgr6*-null = 22.1, P = 0.017) while the number of BSCP was comparable between the genotypes. We also evaluated widely used mesenchymal progenitors marked by Sca1, CD140a, and CD105, to identify mixed populations of mesenchymal progenitor cells. *Lgr6-*null derived bone marrow had fewer CD140a+ (CD140a+, Control = 3.22%, SD ±1.019; *Lgr6*-null =1.54 SD ± 0.47, P =0.012) and CD105+ cells (CD105+, Control = 20.87%; *Lgr6*-null =14.6, P=0.003). *Lgr6-*null periosteal cells also showed a decrease in the expression of CD140a+ cells (CD140, Control = 9.14%; *Lgr6*-null 5.25%, P =0.039). We observed no changes in CD105+ cells in the periosteum. The frequency of Sca1+ cells was comparable between the genotypes in both compartments (Fig. 5e and data not shown).

**Figure 5.**
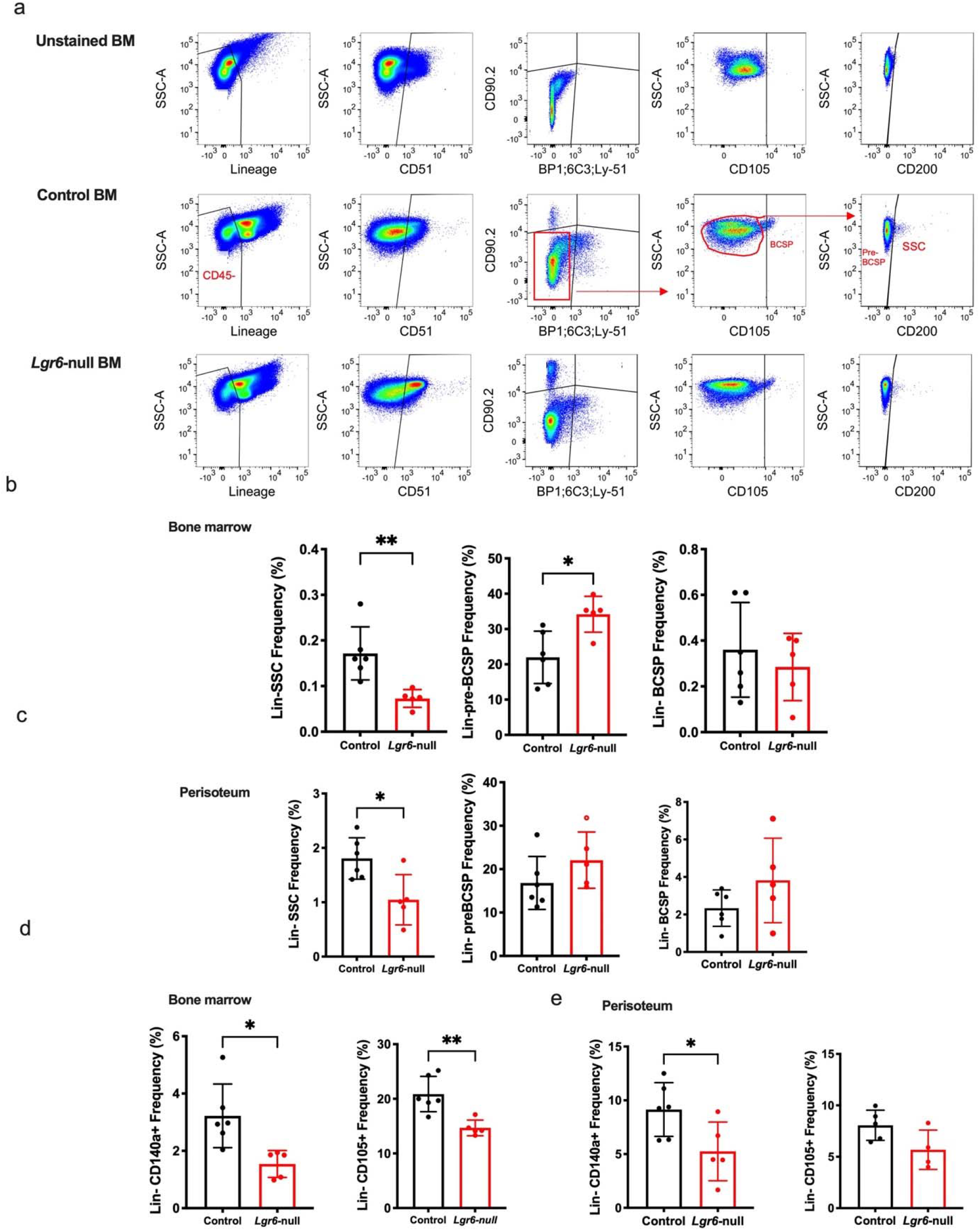
Bone marrow and periosteal cells derived from *Lgr6-*null mice are composed of a different proportion of skeletal stem cells and progenitors. **a**. Gating strategy to determineproportions of skeletal stem cells and their downstream progenitor in bone marrow from unstained, control, and *Lgr6-*null mice. **b**. Bone marrow analysis of SSCs, pre-BSCPs, and BSCP. Graphs represent one experiment using 5-6 male mice/genotype. **c**. Analysis for SSCs, pre-BSCPs, and BSCP population from periosteum-derived cells. For periosteal samples total of 11-12 male mice from each genotype were used, and cells from 2 mice were pooled for each sample. **d and e**. Cells were also analyzed for a specific population of mesenchymal progenitors, Lin^-^CD140^+^ and Lin^-^CD105^+,^ from Bone marrow (d) and periosteum (e). SSC (stem cells), pre-bone cartilage stromal progenitor (pre-BSCP), bone cartilage–stromal progenitor (BSCP). Comparison of control and *Lgr6-*null groups by two-sided Student’s t-test adjusted for non-normality (Mann Whitney test) or unequal variance (Welch’s test) where appropriate. All data are mean SD ±, *p<0.05, **p <0.01.A representative of 3 experiments is shown.

### Lgr6-null mice show deficient proliferation and ALP activity immediately after fracture injury

To determine a functional role for *Lgr6* expression following injury *in vivo*, a closed, stabilized, femoral fracture model of bone healing, an established method to evaluate endochondral regeneration, was used ^10,28,30,40^. To avoid the influence of the reduced bone formation evident at 5 months of age, we employed 3-month-old male and female mice, which do not have any overt skeletal phenotype (data not shown). Upon initiation of fracture repair, skeletal progenitors are activated at the fracture site, and periosteal cells respond to injury within 24-48 h post-injury ^10,30,41^. At 3 days post-fracture, relatively fewer EdU^+^ cells were observed in the periosteum of fractured femurs in Lgr6 null mice compared to controls (EdU^+^ cells, Controls 24%, *Lgr6*-null 13%, P=0.0023) (Fig. 6a). We also analyzed alkaline phosphatase activity; In intact contralateral femurs, level of ALP activity in the periosteum was comparable between the genotypes. However, 3- and-5-days post fracture *Lgr6*-null mice showed respectively a 60% and 48% decrease in the ALP^+^ activity relative to control mice (Fig. 6b). Flow cytometric analysis of mesenchymal progenitors in the periosteum derived from fractured femurs from *Lgr6*-null mice showed 48% decrease in SSCs (P=0.048) and 32% increase in pre-BCSP (P=0.030) populations. In comparison, no changes were observed in the BSCP population (Fig. 6c).

**Figure 6.**
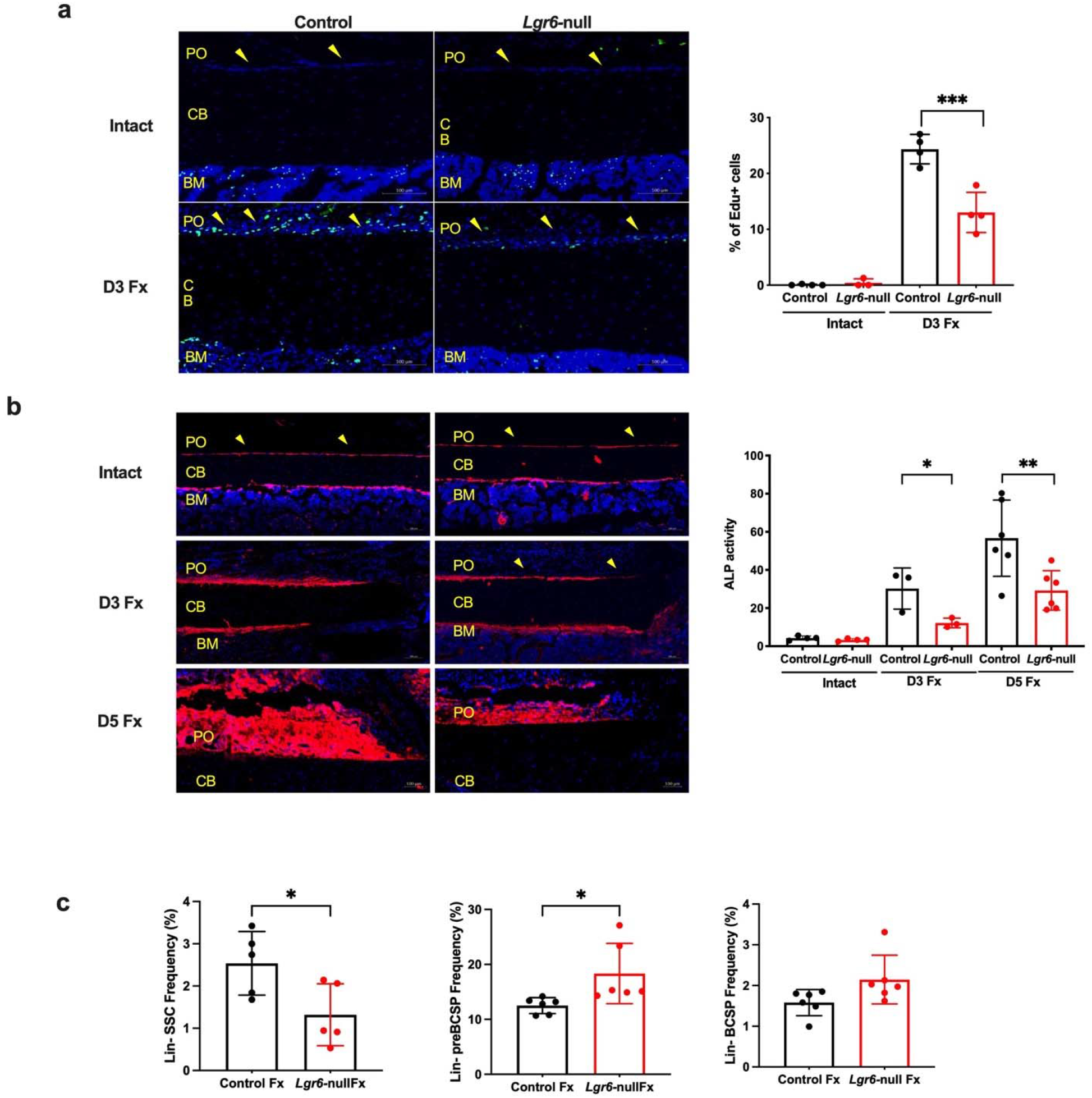
In response to fracture, *Lgr6-*null periosteum is less proliferative and has decreased ALP activity **a**. 10x images of mid-diaphyseal regions in EdU-labeled and DAPI-stained frozen sections from intact, and 3-day fracture femurs from control and *Lgr6-*null mice injected with EdU. Bar graph representing the relative EdU^+^ and DAPI^+^ cells within the periosteal region of control and *Lgr6*-null D3 fractured femurs. n=3-4 male mice/genotype. **b**. *Lgr6-*null mice have decreased ALP activity. 10x images of mid-diaphyseal regions in ALP and DAPI-stained frozen sections from intact, 3 -and 5-days post-fracture femurs from control and *Lgr6-*null mice. Bar graph representing the percent ALP^+^ periosteal pixels over a fixed threshold of 85 normalized to the total pixels per ROI. n=3-6 male mice/genotype. PO: periosteum, CB: Cortical bone, BM: Bone marrow. **c**. Periosteal cells isolated from the fracture site were analyzed for a population of mouse skeletal stem cells defined as CD90-CD200+ cells from the parent Lin-CD51+Ly51-CD105-population and the downstream bone, cartilage, and stromal progenitors (BCSPs), defined as CD90-CD105-cells from the parent Lin-CD51+Ly51-population. Cells from 2-3 fractured femurs were pooled, and 11-12 male mice for each genotype were used. A representative of 2experimentst is shown. All data are mean SD ±. *p<0.05, **p <0.01.

### *Lgr6-*null mice have impaired endochondral fracture healing

Following femoral fracture in *Lgr6-*null mice and controls, we analyzed the calluses during the bone regeneration phase (14 days post-fracture) and remodeling phase (28 days post-fracture) to evaluate healing. While overall callus size at 14 days post-fracture was comparable among groups, *Lgr6-*null calluses had significantly reduced bone content/volume relative to controls. Reconstructed images from μCT scanning demonstrated the presence of large unmineralized voids in *Lgr6-*null fracture calluses (Fig. 7a). μCT analysis showed that *Lgr6*-null mice had significantly reduced callus BV/TV as compared to control (BV/TV, Control = 22.3 %; *Lgr6*-null = 15.47%, P=0.019). Additional analysis revealed a 22% decrease in bone mineral density (BMD) in *Lgr6*-null fracture calluses compared to controls (Fig. 7b). Safranin O staining and cellular morphology indicated that these voids were composed of cartilaginous tissue and that *Lgr6-*null calluses had retained 41% more cartilage content compared to the controls at this phase of fracture repair, lacking the robust mineralization capacity of control samples (Fig. 7c and d).

**Figure 7.**
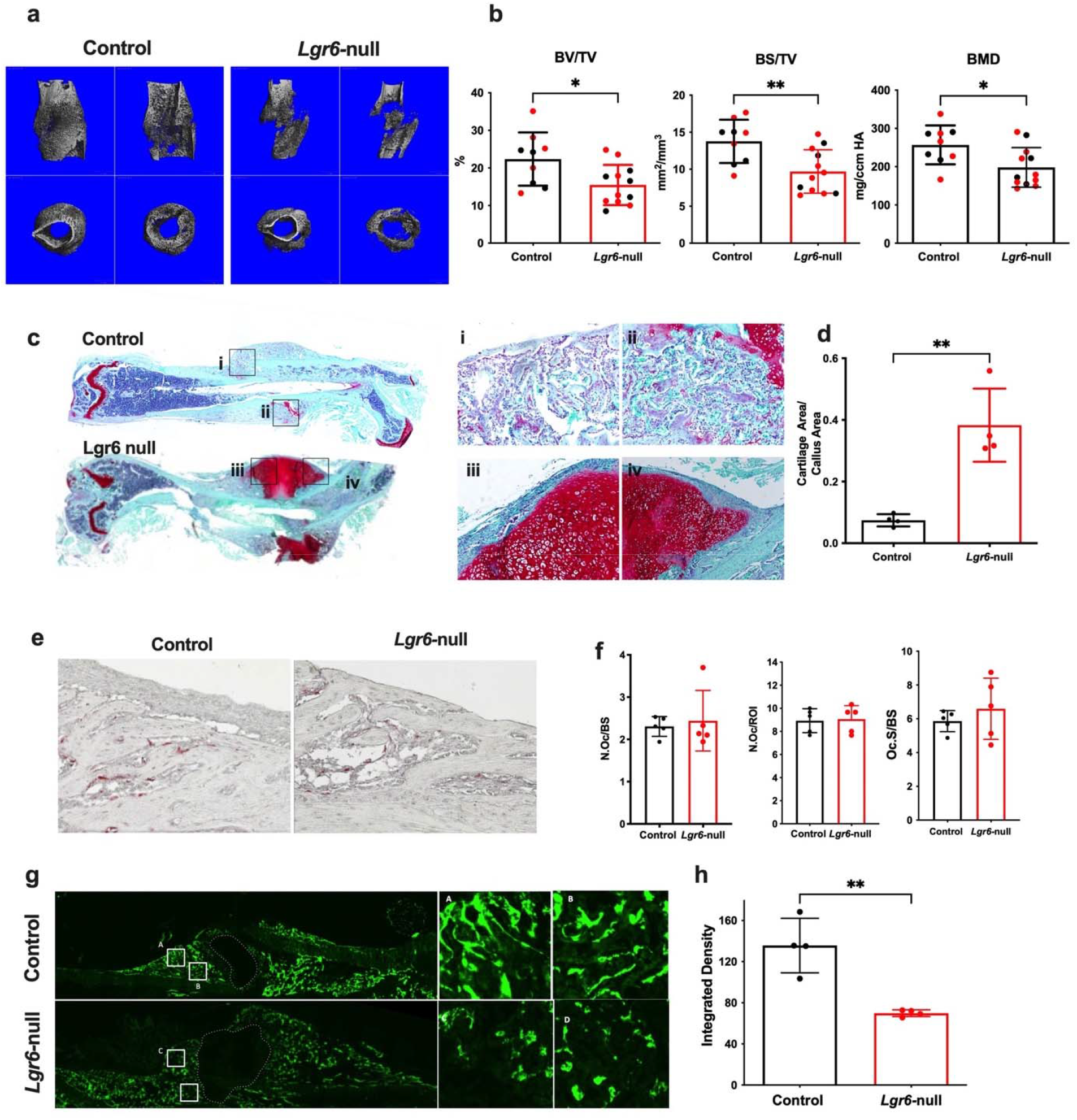
*Lgr6-*null mice experience impaired endochondral fracture healing. **a**. Representative 3D μCT images of calluses from fractured femurs 14 days after fracture. **b**. μCT parameters of calluses 14 days post-fracture in control animals compared to *Lgr6*-null. BV/TV=bone volume per total volume within the callus TV=total callus volume. 12-week-old males (red dots) and females (black dots). n=9-12 mice/genotype **c**. Histological section stained with Safranin O/fast green stains of representative fractured bones at day 14 post-fracture. **d**. Quantification of cartilage area in the callus. n=4 mice/genotype. **e**. TRAP staining **f**. quantification of osteoclast parameters from areas flanking the fracture site 14 days post-fracture. n=5 mice/genotype. Scale bar=100μm. N. Oc/ROI=number of osteoclasts in the region of interest, N. Oc/BS=number of osteoclasts per bone surface, Oc. S/BS =osteoclast surface per bone surface. **g**. Calcein was injected one day before sacrifice to label areas of active bone formation in mineralizing areas of the healing fracture. Fluorescent microscopy images demonstrating the different calcein labeling patterns between and control fracture calluses, with fluorescence integrated density quantified in (h). n=4/male mice per genotype. Scale bar=1000μm.

During the regeneration phase, in areas flanking the central fracture callus, we observed successful new bone formation in the *Lgr6-*null samples in histological sections and via μCT. TRAP staining and quantification showed no differences in the number of osteoclasts or osteoclast surfaces compared to the total bone surface in these areas of ossification flanking the fracture site (Fig. 7e and f). To indirectly evaluate the *in vivo* osteogenic activity of osteoblasts in these areas of the calluses, we injected calcein one day before animal sacrifice to label the active bone surfaces (Fig. 7g and h). We found that even in areas of mineralization within the *Lgr6-*null calluses, there was reduced osteoblast activity compared to controls, as evident from the mineralization pattern and the quantified integrated density of calcein fluorescence within these defined regions of interest (Control =186; *Lgr6*-null = 101, P =0.025).

We then evaluated the remodeling phase of fracture healing to observe the calluses’ ability to ossify and eventually remodel. We found that the *Lgr6-*null calluses ultimately mineralized successfully (Fig. 8a). However, *Lgr6-*null calluses had 25% reduced bone volume fraction and 23% decreased bone mineral density compared to controls, as measured by μCT (Fig. 8b). Additionally, the healing bones of *Lgr6-*null mice continued to display aberrant morphology compared to controls, with significantly larger calluses in terms of total volume (TV mm^3^ Control= 7.98 ± 4.73; *Lgr6*-null = 22.1 ± 6.98, P=<0.0001), but the very sparse formation of new trabeculae. Safranin O staining confirmed the complete lack of cartilaginous matrix within the calluses, and a counterstain with fast green showed fewer and thinner areas of newly formed bone in *Lgr6*-null-derived samples compared to controls, corroborating with μCT data (Fig. 8c). TRAP staining of defined regions within these calluses revealed significantly more osteoclasts per bone surface (N.Oc./BS, Control = 6.3; *Lgr6*-null = 10.9, P= 0.005) present in the day 28 post-fracture *Lgr6-*null calluses (Fig. 8d). A calcein label showed increased osteoblast activity (Control = 70; *Lgr6*-null = 119, P= 0.0002) in the *Lgr6-*null calluses compared to control (Fig. 8e). These results combined demonstrated higher bone remodeling activity in *Lgr6-*null calluses, while control calluses remodeled and closer to complete healing, overall indicating delayed healing progression.

**Figure 8.**
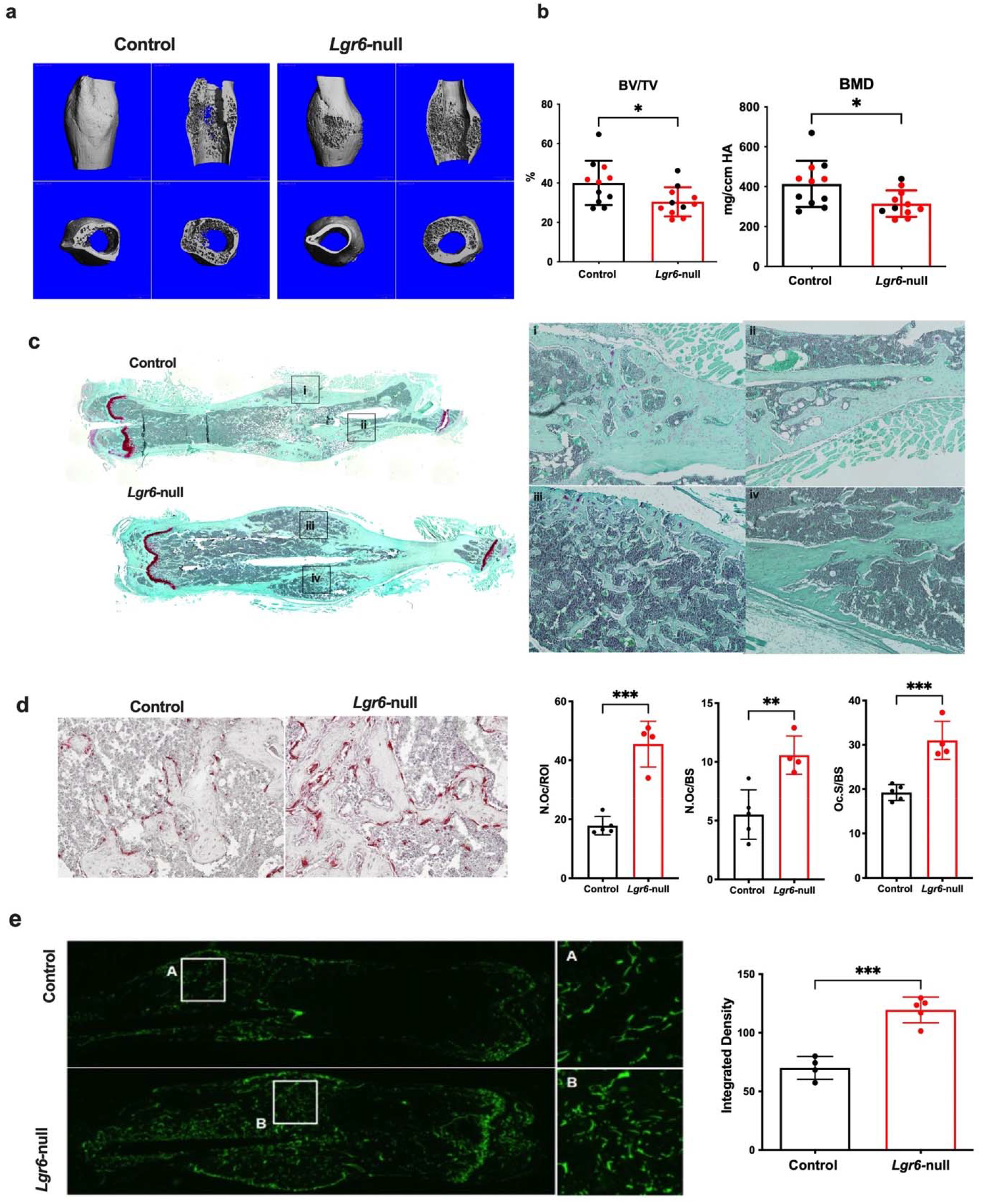
Post-fracture bone remodeling is delayed in *Lgr6*-null mice. **a**. Representative 3D μCT images of calluses from fractured femurs 28 days after fracture, and b. corresponding μCT analysis of fractured bones. 10-12-week-old males/genotype (red dots) and females (black dots). n= 8-11 mice/genotype. **c**. Safranin O/fast green staining of control and *Lgr6-null femurs* 28 days post-fracture. **d**. TRAP staining **e**. quantification of osteoclast parameters from areas flanking the fracture site 28 days post-fracture. Scale bar=100μm. N. Oc/ROI=number of osteoclasts in the region of interest, N. Oc/BS=number of osteoclasts per bone surface, Oc. S/BS=osteoclast surface per bone surface. Calcein was injected one day before sacrifice to label areas of active bone formation in mineralizing areas of the healing fracture. **a**. Fluorescent microscopy images demonstrating the different calcein labeling patterns between *Lgr6-*null and control fracture calluses 28 days after fracture. Fluorescence integrated density quantified in (**b**). n=3-5 male mice/genotype.

## Discussion

In this study, we demonstrate that Lgr6 is distinct among the three Lgr family members found in mammals, as it is associated with regeneration and osteogenic capacity rather than skeletal development. Single-cell RNA-sequencing and microarray analysis of long bones showed relatively high expression of Lgr6 on the skeletal stem compared to other skeletal stem/progenitor cells regulated by cWnt signaling. Furthermore, in response to bone injury, the number of Lgr6+ cells and their progeny was increased at the site of new bone formation. Despite representing a minor fraction of the total population, this gene is necessary to mediate proper bone regeneration since the loss of *Lgr6* expression results in impaired endochondral ossification during fracture healing.

Importantly, in strong support of our work examining the role of Lgr6 in fracture healing, a recent RNA expression analysis of the mesenchymal stem cells derived from fractured bones of patients with hip fractures found that *LGR6* was the most upregulated gene (Fold change 11.0, corrected P-value =0.0083) ^42^. Furthermore, separate GWAS studies have shown that variants associated with *LGR4* and *LGR6* loci are correlated with low BMD, osteoporosis risk, and osteoporotic fractures ^43,44^. In addition, human genetic studies showed that increased *RSPO3* expression is associated with increased trabecular bone mineral density and reduced risk for fractures ^45^. Combined these studies underscore the role of Rspo/Lgr axis in bone regeneration.

cWnt signaling is activated during fracture healing and bone regeneration; osteoblast-specific β-catenin null mice show severely inhibited fracture repair ^46^, and loss of β-catenin signaling in chondrocytes reduces fracture healing capabilities ^47^. Bmp2 functions downstream of Wnt in the periosteal layer, a critical reservoir of skeletal progenitors, and activates *Sp7* transcription within regenerating bone ^48^. Loss of *Bmp2* in cells that have already passed the *Sp7*+ “checkpoint” does not affect Wnt signaling later in differentiation, suggesting that uncommitted osteoblast progenitors are a putative target of *Bmp2* ^*49*^. These studies shed light on our results in both *in vivo* fracture studies and *in vitro* differentiation assays of mesenchymal progenitors, as Lgr6 may be involved in upstream Wnt/Bmp2-mediated endochondral fracture healing. Further studies are necessary to understand whether, for endochondral repair, Lgr6 is specifically required in bone regeneration to enhance or modulate signaling by either cWnt or BMP2 or both.

We found that the regeneration phenotype in *Lgr6-*null mice is due to a decrease in cell proliferation during the initial phase of impaired fracture healing and a defect in osteogenic capacity and osteoblast differentiation. However, there was no detrimental effect on the cartilage-to-bone transition ^50,51^, as calluses of *Lgr6-*null fractured bones eventually ossify successfully. This finding contrasts with other fracture healing studies reporting the impairment of successful cartilage differentiation during bone healing, for example, in a mouse model with an over-activating mutation of *Fgfr3* in Prx1-derived progenitors ^52^. In this study, periosteal cells with the *Fgfr3* mutation do not undergo expected terminal chondrocyte hypertrophy following transplantation^53,54^, leading to failure of fractures to consolidate; the chondrocytes fail to hypertrophy and terminally differentiate, and there is persistent fibrosis within the callus day 28 post-fracture. In our study, calluses from *Lgr6-*null animals at day 28 post-fracture are still being heavily remodeled and have less mineralized tissue content due to delays in the healing process; however, there is no evidence of fibrosis or cartilage remnants at this time point.

Using lineage tracing and flow cytometric approaches, several groups have identified distinct skeletal stem and progenitor cell populations ^39,55-57^ in complex niches ^58-61;^ however, some of these phenotypic markers are also expressed on mature osteoblasts^62^. Our flow analysis indicates that bone marrow and periosteum in *Lgr6-*null mice have a smaller proportion of SSC than controls, but there is an increase in the pre-BSCP population. One possible mechanism may be that since *Lgr6* is a direct target of Wnt signaling, in the context of cWnt/BMP2 interplay for modulation osteogenic differentiation, Lgr6 likely regulates the transition of Runx2/Osterix+ cells to a more differentiated state. In support of this notion, treating BMP2-Prx-1 null cells with Wnt 3a decreased Lgr6 expression (p<0.003) ^49^. Another possibility is that Lgr6-responsive stem/progenitor cells control “stemness” by maintaining the expression of inhibitors of differentiation. While the idea that Lgr6 signaling contributes to the maintenance of immature mesenchymal progenitor cells through the induction of an anti-differentiation program is intriguing, further characterization of this pathway in mesenchymal progenitors is needed.

In Lgr6-null cultures, we found no changes in osteoclast differentiation but demonstrated decreased ALP+ colonies, osteogenic differentiation, and mineralization; therefore, we attribute the decreased trabecular bone volume in *Lgr6*-null mice to the reduced bone formation by osteoblasts. The Lgr6-null trabecular phenotype is more pronounced than the cortical phenotype. Other models of deficient Wnt signaling, including Wnt10B ^63^, Wnt16 ^64-66^, Notum ^67^, Rspo2 ^68^, and Rspo3 ^45^, also have shown a discrepancy between trabecular and cortical bones. Further studies in *Lgr6*-null mice at geriatric time points are needed to examine the effect of Lgr6 on cortical bones. In agreement with our studies, another global Lgr6 knockout mice with a deletion in exons 15 and 16 showed a decrease in bone volume in both sexes compared to control cohorts at 2 months of age, but no changes were found in cortical bone parameters ^19^. In our studies, mice lacking endogenous *Lgr6* expression at 3 months of age had no overt skeletal phenotype, likely due to the compensatory effect by other Lgr family members ^69,70^; however, with age, this compensation is insufficient in maintaining bone volume. Further, this compensatory mechanism is not evident upon injury since fracture healing is significantly impaired in 3-month-old *Lgr6*-null mice. Together, these data support the notion that Lgr6+ osteoprogenitors are activated in response to bone injury.

The effect of Wnt signaling on regulating osteoprogenitors is complex: the disruption of cWnt signaling decreases progenitor number in the Lrp5-knockout, Wnt10b-knockout, Rspo2 knock out models ^63,68,71^ but high levels of expression of stabilized beta-catenin also reduced the number of CFU-F colonies ^72^. While no difference in Axin2 or β-catenin expression (data not shown) was observed in long bones of *Lgr6*-null mice, *in vitro* studies showed that in *Lgr6*-null cells in response to Wnt3a stimulation, expression of Wnt response genes, *Axin2* and *Lef1* was attenuated. However, the levels of these proteins also depend on the osteogenic differentiation; therefore, a direct comparison of Wnt signalosome components between *Lgr6-*null cells and controls following osteogenic induction may still not lead to a definitive answer, as *Lgr6-*null cells have impaired osteogenic differentiation *in vitro*. Thus, it will be difficult to interpret if the results are due to alterations in Wnt signaling or osteogenic differentiation potential by another mechanism unassociated with Wnt. Nevertheless, it cannot be ruled out that *Lgr6* acts in a currently unidentified, Wnt-independent mechanism, especially considering recent reports on Lgr family member involvement in different signaling pathways, including ERK/FGF, BMP2, and BMP9 ^36,73,74^.

Currently, known ligands for Lgr6 include R-spondins and Maresin 1 (MaR1). MaR1 delivery to tibial fractures enhanced bone healing ^78^, and while MaR1 can bind Lgr6 ^79,80^, studies are needed to demonstrate a MaR1/Lgr6 interaction specifically within osteoblast lineage. Rspo2 is an essential regulator of bone mass ^68^, and various Rspos may enhance osteogenic differentiation *in vitro* ^*75-77*^. However, Rspos have binding affinity for all three Lgr family members, and it remains unclear if a distinct Rspo/Lgr6 axis regulates osteoblast differentiation^31^. Further, Rspo2 is a possible ligand within the BMP pathway, which may be the mechanism by which it enhances osteogenesis^73^.

Our lineage tracing studies have some limitations. Despite the inclusion of a GFP reporter in the Lgr6^EGFP-ires-CreERT2^ model, a direct comparison between *Lgr6*-positive progenitors and *Lgr6*-negative progenitors was not performed. Since we were unable to isolate GFP+ cells due to a very weak signal and a lack of available antibodies appropriate for use in these applications. The Lgr6^EGFP-ires-CreERT2^ model is a low-efficiency Cre mouse model; we found that under homeostatic/non-injured conditions, tamoxifen-induced fewer numbers of Lgr*6*+ cells in the bone compartment. While there are significantly increased numbers of Td tomato+ cells in fracture callus, a new mouse model to use as a lineage tracing tool that more faithfully marks Lgr6+ cells upon tamoxifen induction would be informative.

In summary, our studies demonstrate that *Lgr6-*null mice experience significant delays in callus mineralization and healing during endochondral fracture repair. The osteogenic differentiation and *in vivo* regeneration phenotypes in *Lgr6-*null mice indicate that this gene has an essential role in postnatal osteoblastogenesis and skeletal regeneration. The dynamic expression of *Lgr6* during osteogenic differentiation and its important role in endochondral fracture repair warrant further studies to understand a complete mechanism of skeletal regeneration.

## Methods

### Mice

*Cbl*^*YF/YF*^ mice were described previously ^25^. Mice with knock-in targeted insertion of the EGFP-Ires-CreERT2 into the Lgr6 transcriptional start of *Lgr6* locus (*Lgr6*^*EGFP-IRES-CreERT2*^) were bred into a mixed 129/SV background^16^. Heterozygous breeding of these knock-in mice into homozygotes yields *Lgr6*^*EGFP-IRES-CreERT2/*^ *Lgr6*^*EGFP-IRES-CreERT2*^ (henceforth *Lgr6*-null mice), *Lgr6*^*EGFP-IRES-CreERT2*^*/*^*+*^ and control littermates; functional endogenous *Lgr6* expression is blocked from both alleles in *Lgr6*-null mice^16,81^. *Lgr6*-null and *Lgr6*^*EGFP-IRES-CreERT2*^ */*^*+*^ mice developed normally with no overt developmental phenotype^32,81^. In *Lgr6*^*EGFP-ires-creERT2*^*/*^*+*^ mice, due to extremely weak IRES reporter, we do not see GFP+ cells in either bone sections or bone cells by histology and flow cytometry (data not shown). In our studies, we compared *Lgr6*-null and control mice since we found no difference in the skeletal phenotype of control and *Lgr6*^*EGFP-IRES-CreERT2*^ */*^*+*^ mice of either sex at 3- or-5 months of age (data not shown). For lineage tracing experiment, *Lgr6*^*EGFP-IRES-CreERT2*^ */*^*+*^ were bred with Ai9 tomato reporter (Ai9 (B6. Cg-Gt (ROSA) 26Sor^tm9(CAG-tdTomato) Hze^/J). Both male and female mice were used as indicated. Genotyping for mice was performed at the time of weaning and before each experiment using Lgr6-specific primers listed in Supplementary Table 1.

### Animal Welfare

This study was approved by the local IACUC and was conducted per the national legislation on the protection of animals and the NIH Guidelines for the Care and Use of Laboratory Animals. All animals were housed in ventilated cages in a temperature- and humidity-controlled environment under a 12-hour light cycle.

### μ*CT analysis*

Structural analysis of intact and fractured femurs was performed using cone-beam x-ray microcomputed tomography (μCT40, Scanco Medical AG) at the μCT Imaging Facility at UConn Health. 8μm^3^ scans were acquired at 55kVp, 145μA, 1000 projections with a Gaussian filter. Intact femurs were analyzed at the microCT Core at UConn Health, with lower thresholds of 270 per mille (449.4mgHA/ccm) for trabecular bone and 400 per mille (748.2mgHA/ccm) for cortical bone. Fractured femurs were analyzed by manually segmenting callus tissue from surrounding bone^82^, with a lower threshold of 270 per mille (403.3mgHA/ccm).

### Cell isolation

Bone marrow and periosteal cells were isolated from femurs and tibias as previously described^30^. For periosteal cultures, 2-3 mice of the same sex and genotype were pooled to get enough cells. Briefly, the periosteum was scraped from flushed, hollow bones into sterile PBS and digested enzymatically (0.05% collagenase P, 0.2% hyaluronidase in PBS) for one hour at 37° C in an orbital shaker. Then, complete media was added to stop enzymatic digestion, and cells were filtered through a 40 μm mesh strainer to form a single-cell suspension and remove debris. Cells were grown in 5% oxygen for 4 days to allow for expansion before transfer to 20% oxygen.

### CFU Assays

Colony-forming unit (CFU) assays were performed using bone marrow-derived and periosteum-derived cells. Cells were isolated as described above. Bone marrow-derived cells from individual mice were directly plated at 1×10^6^ cells 60 mm culture plates in duplicates and cultured for 10 days in 20% oxygen without cell passage. Periosteal cells were pooled from two mice of the same sex, age, and genotype for each biological replicate for appropriate cell numbers. Cells were plated at 5×10^5^ cells per well in 60 mm plates in duplicates and cultured for 4 days in 5% oxygen, followed by 4 days in 20% oxygen without cell passage. Fixed cells were stained either for ALP activity using an ALP detection kit according to the manufacturer’s instructions (Sigma-Aldrich) or with crystal violet. Colonies were imaged using a Leica microscope and were evaluated using converted greyscale 8-bit images in ImageJ

### Cell differentiation and staining

For osteogenic differentiation, periosteal- and bone marrow-derived cells were seeded (2×10^5^ cells/well) in 12-well plates. Cells were treated with differentiation media (50 μg/mL ascorbic acid, 4 mM β-glycerophosphate) once cells reached 80-90% confluence in culture in about 2-3 days. Media was changed every other day for 7, 14, and 21 days. Cultures were stained to detect ALP activity as described above. Cells were fixed in cold methanol and stained with Alizarin Red S (Science Cell) to visualize *in vitro* mineralization. Staining was quantified according to the manufacturer’s instructions using colorimetric detection. Osteogenic differentiation was repeated at least three times per time point to confirm ossification trends. For chondrogenic assays, periosteal bone marrow-derived cells were seeded in 12-well plates were treated with chondrogenic differentiation media starting when cells reached 80-90% confluence, as we have previously described^30^. Media was replaced every other day for the remainder of the assay. After 14 days, cells were either harvested to isolate RNA or were fixed and stained with Alcian blue to assess chondrocytic differentiation. Osteoclast cultures were established and evaluated as previously described^29^.

### Bulk mRNA sequencing and analysis

RNA was isolated from cell cultures using a Trizol (Invitrogen) extraction method. Total RNA was quantified, and purity ratios were determined for each sample using the NanoDrop 2000 spectrophotometer (Thermo Fisher Scientific). To further assess RNA quality, total RNA was analyzed on the Agilent TapeStation 4200 (Agilent Technologies) using the RNA High Sensitivity assay. Only samples with RINe values above 8.0 were considered for library preparation. Illumina Transcriptome Library preparation and Sequencing Total RNA samples were prepared for mRNA-Sequencing using the Illumina Stranded mRNA Ligation Sample Preparation kit following the manufacturer’s protocol (Illumina). Libraries were validated for length, and adapter dimer removal using the Agilent TapeStation 4200 D1000 High Sensitivity assay (Agilent Technologies), then quantified and normalized using the dsDNA High Sensitivity Assay for Qubit 3.0 (Life Technologies). Sample libraries were prepared for Illumina sequencing by denaturing and diluting the libraries per the manufacturer’s protocol (Illumina). All samples were pooled into one sequencing pool, equally normalized, and run as one sample pool across the Illumina NovaSeq 6000 using version 1.5 chemistry. A target read depth of 50 M reads was achieved per sample paired-ended end 100bp reads. Raw reads were trimmed with FASTP 96 (version 0.23.0), with low-quality sequences removed, and trimmed reads mapped to Mus 97 musculus genome (GRCm39) with HISAT2 (version 2.1.1). SAM files were then converted into BAM 98 format using samtools (version 1.12), and the PCR duplicates were removed using PICARD 99 software (http://broadinstitute.github.io/picard/). Counts were generated against the features with 100 htseq-count. Differential expression of genes between conditions was evaluated using DESeq2 101.

### RNA isolation and cDNA synthesis for qPCR

Total RNA was extracted from cultured cells and freshly dissected tibia. For tibial RNA extraction, both ends of each tibia were cut off to remove the growth plates, and the marrow was flushed by centrifugation at 300 g for 1 minute at 4°C. Both tibiae from one mouse were pooled and homogenized in TRIzol (Invitrogen). Tissue RNA integrity was checked on BioAnalyzer RNA NanoChips (Agilent Technologies) at the UConn Health molecular core facility. Only samples with a RIN value of <8.0 were used. cDNA synthesized using Superscript III cDNA synthesis kit (Invitrogen). qRT-PCR reactions were performed on a Bio-Rad real-time PCR system using SYBR Green (BioRad) PCR reagents. For each gene of interest, qPCR was performed in both technical and biological triplicates. Data were normalized to GAPDH or beta-actin. The 2^-ΔΔCt^ method was used to calculate the relative gene expression. Primer sequences are listed in Supplementary Table 1.

### Assessment of cWnt Signaling

To analyze the expression of genes by real-time PCR, bone marrow-derived stromal cells (BMSC) were cultured to confluence, serum starved for 12h, and then treated with Wnt3a (50ng/ml) (R and D systems). Seventy-two hours after treatment, cells were lysed with Trizol (Life Technologies) and frozen at −80° C. Subsequently, RNA was extracted, and qRT-PCR was performed using Axin2 and Lef1 primers listed in Supplementary Table 1.

### Flow Cytometry

Bone marrow cells were isolated from each mouse as one biological replicate. For periosteum-derived preparations, cells from 2-3 mice were pooled. For fractured samples, 2-3 fractured femurs were pooled. In all cases, after cell isolation, red blood cell lysis was performed in ACK buffer, as described above. For each sample, 1×10^6^ cells were collected in cell staining media containing HEPES and 2% heat-inactivated FBS in Hanks Balanced Salt Solution (HBSS). Cells were incubated with antibodies and secondary reagents indicated in Supplementary Table 2. Cells were incubated with an antibody cocktail for 45 minutes on ice and then washed in ice-cold staining media. Dead cells were excluded using Live IR staining. Cells were analyzed using a BD-LSRII flow cytometer (BD Biosciences). All experiments included unstained bone marrow and periosteal cells for establishing gates. Fluorochrome beads were used as single-color controls. In some experiments, fluorescence minus one (FMO) control was used to assist in gating. Experiments included 5-6 biological replicates and were performed twice to ensure consistent outcomes.

### Surgical Procedures

Closed, stabilized femoral fractures were generated as previously described in 10-week-old mice^28,40^. Briefly, pin placement was confirmed using a Faxitron Cabinet X-Ray system (Faxitron Corporation), and the femur was fractured using an Einhorn three-point drop-weight device and evaluated using X-ray. Buprenorphine (0.1 mg/kg body weight) was administered by subcutaneous injection before the procedure and twice a day for three days following the fracture for pain management. Samples excluded from the study analysis included multi-fracture, bent pins, and fracture placement that was not mid-diaphyseal. Both male and female mice were used in fracture healing experiments, and mice were euthanized 3, 5, 1428 days8-days post-fracture for analysis.

### Lineage Tracing

To activate the inducible Cre in the Lgr6^EGFP-ires-CreERT2^/^+^; Ai9/^+^ mouse model, tamoxifen (10 mg/mL, Sigma-Aldrich) in corn oil or vehicle was administered intraperitoneally at a dosage of 75 μg/g body weight in 10–12-week-old-mice, and long bones were harvested 5 and 14 days after injection. In some experiments, tamoxifen was administered one day before and the day of fracture, and mice were sacrificed for histological analyses 5-and 14-days following fracture surgery. Bones were processed for frozen sectioning as previously described^40^.

### Single-Injection Calcein Labeling

For visualizing remodeling fracture calluses, calcein was injected (10 mg/kg body weight IP) one day before sacrifice to label active areas of bone formation, and a small piece of the tail was clipped to visualize the presence of dyes on the day of the injection integrate dated density of the calcein fluorescent label was quantified in ImageJ. Measurements were taken from four different sites of intramembranous bone formation, flanking the fracture site of each sample, and averaged.

### Assessment of cell proliferation and ALP activity in response to fracture

Mice were injected with 5-ethynyl-2’-deoxyuridine (EdU) (10mM) in normal saline 24 hours before sacrifice to measure cell proliferation. Fractured and intact contra lateral femur sections were stained for EdU labeling using the Click-iT® EdU Alexa Fluor® 488 Imaging Kit (Life Technologies) according to the manufacturer’s instructions. ALP activity post-fracture was evaluated using a modification of the ALP Leukocyte kit (86R-1KT) (Sigma) with Fast Blue-BB salt (F3378). ALP^+^ periosteal area was quantified using methods described previously ^30^. RGB images were split into the three-color channel 8-bit images. The red channel 8-bit image was analyzed further for positive pixels in the region of interest (ROI) in the periosteum, above the threshold of 55 for intact femurs and 85 for D3 fractured femurs. Data are presented as a percentage of ALP^+^ positive pixels over the threshold, normalized to total pixels in the ROI measured. To visualize morphology and detect cartilage content, fixed and decalcified samples were paraffin-embedded decalcified bone sections were cut at 7 μm on a microtome, sections were deparaffinized, stained with safranin O/fast green, and counterstained with hematoxylin to detect cartilage content. Tartrate-resistant acid phosphatase (TRAP) staining was performed to visualize osteoclasts in fractured bones. Sections were analyzed using Osteomeasure^30,40^ to measure the number of TRAP^+^ osteoclasts, osteoclast surface, and bone surface in regions of interest.

### Statistical Analyses

Experiments conducted for this study were repeated at least three times or were presented with biological replicates. Statistical analyses were performed using GraphPad Prism v6 software with either a student’s t-test or one-way ANOVA with Tukey’s post hoc test comparing the control group to *Lgr6*-null-mice. Data are presented as mean ± SD. *P-values* < 0.05 were considered statistically significant.

## Acknowledgments

Funding was provided by NIH/NIDCR R01-DE030716-01 and NIH/NIAMS R01-DE030716-01S1 to AS and KDH, NIH/NIAMS R01-AR070813 to IK, and the NIH/NIDCR training grant T90-DE021989 and NIH/NIDCR F30-DE029100 to LD. We recognize the contribution of Dr. Bo Reese and Dr. Vijender Singh from UConn Center For Genome Innovation. We also appreciate the help provided by Ms. Jessica Kraus in staining paraffin sections.

## Contributions

LD and MW generated fractures, performed cryohistology, staining, imaging, characterized cell cultures, performed quantifications, and data analysis. AP performed flow cytometry for intact and fractured bones and analyzed data. DWY performed micro-CT and data analysis on intact and fractured bones. JSK performed computational analysis of published RNA seq data. LD performed the bulk mRNA sequencing and analysis and wrote the first draft of the manuscript. IK and KDH critically read the manuscript. AS conceived the experimental design, wrote the final version of the manuscript, and supervised the project.

## Figure Legends

Supplementary Figure 1. Characterization of *Lgr6-*null mice. **a**. qRT-PCR analyses of Lgr family members. **b**. Expression of Lr6 protein was detected by Western blotting. HeLa cell lysate was used as a positive control. **c**. No difference was observed in the weight or lean mass of the control of Lgr*6-*null mice. **d**. *Col1a1, Runx2, and Axin* expression levels in the tibia were comparable between control and *Lgr6-*null mice.

Supplementary Figure 2. qRT-PCR analyses of critical osteogenic genes on days 14 and 21 of osteogenic differentiation of periosteal cultures. A representative of 4 experiments is shown **b**. Periosteal cells underwent chondrogenic differentiation in vitro and were stained with Alcian blue or subjected to qRT-PCR analyses. **c**. *In vitro* osteoclastogenesis assay of control and *Lgr6*-null-derived cells, stained for TRAP. TRAP+ cells with >3 nuclei were considered osteoclasts, and osteoclasts per well were counted manually. A representative of 2 experiments is shown

---

**Supplementary Table 1.**
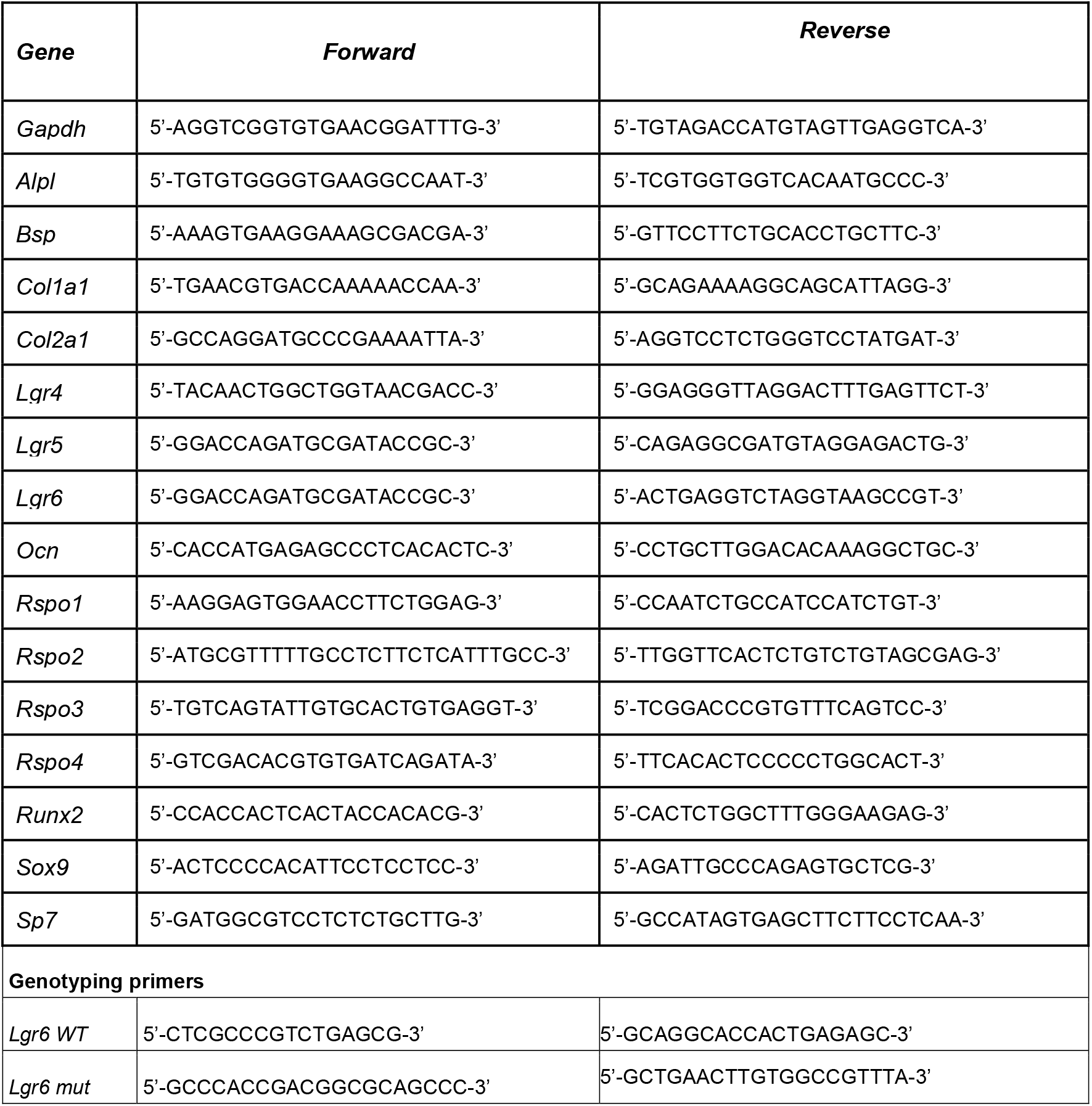

**Supplementary Table 2.**

| Reagent Type | Designation | Source or reference | Identifiers | Additional information |
| --- | --- | --- | --- | --- |
| Antibody | Lineage cocktail Pacific Blue™ anti-mCD3/ Pacific Blue™ anti-mLy-6G(Ly-6C)/ Pacific Blue™ anti-mCD11b/ Pacific Blue™ anti-mCD45R(B220)/ Pacific Blue™ anti-mTer-119 | Biolegend | 133306 | Flow (1:400) |
| Antibody | Pacific Blue™ anti-mouse CD31 Antibody | Biolegend | 102422 | Flow (1:400) |
| Antibody | Brilliant Violet 510™ anti-mouse CD90.2 (Thy-1.2) Antibody | Biolegend | 140319 | Flow (1:400) |
| Antibody | Ly-6A/E (Sca-1) Monoclonal Antibody (D7), Alexa Fluor™ 700, eBioscience™ | eBioscience™, Thermo Fisher Scientific | 56-5981-82 | Flow (1:400) |
| Antibody | PE/Cyanine7 anti-mouse Ly-51 Antibody | Biolegend | 108314 | Flow (1:200) |
| Antibody | APC anti-mouse CD200 (OX2) Antibody | Biolegend | 123810 | Flow (1:200) |
| Antibody | CD140a (PDGFRA) Monoclonal Antibody (APA5), PerCP-eFluor 710, eBioscience™ | eBioscience™, Thermo Fisher Scientific | 46-1401-82 | Flow (1:400) |
| Antibody | BV650 Rat Anti-Mouse CD51 | BD Biosciences | 740546 | Flow (1:100) |
| Antibody | BV711 Rat Anti-Mouse CD105 | BD Biosciences | 740808 | Flow (1:200) |
| Nucleic Acid Stain | Propidium iodide solution (P3566) | Invitrogen, Thermo Fisher Scientific | P3566 | Flow (1:500) 3 µM final, 1.5 mM stock solution |

