## Supplemental Figures for "Wnt-associated adult stem cell marker Lgr6 is required for osteogenesis and fracture healing"

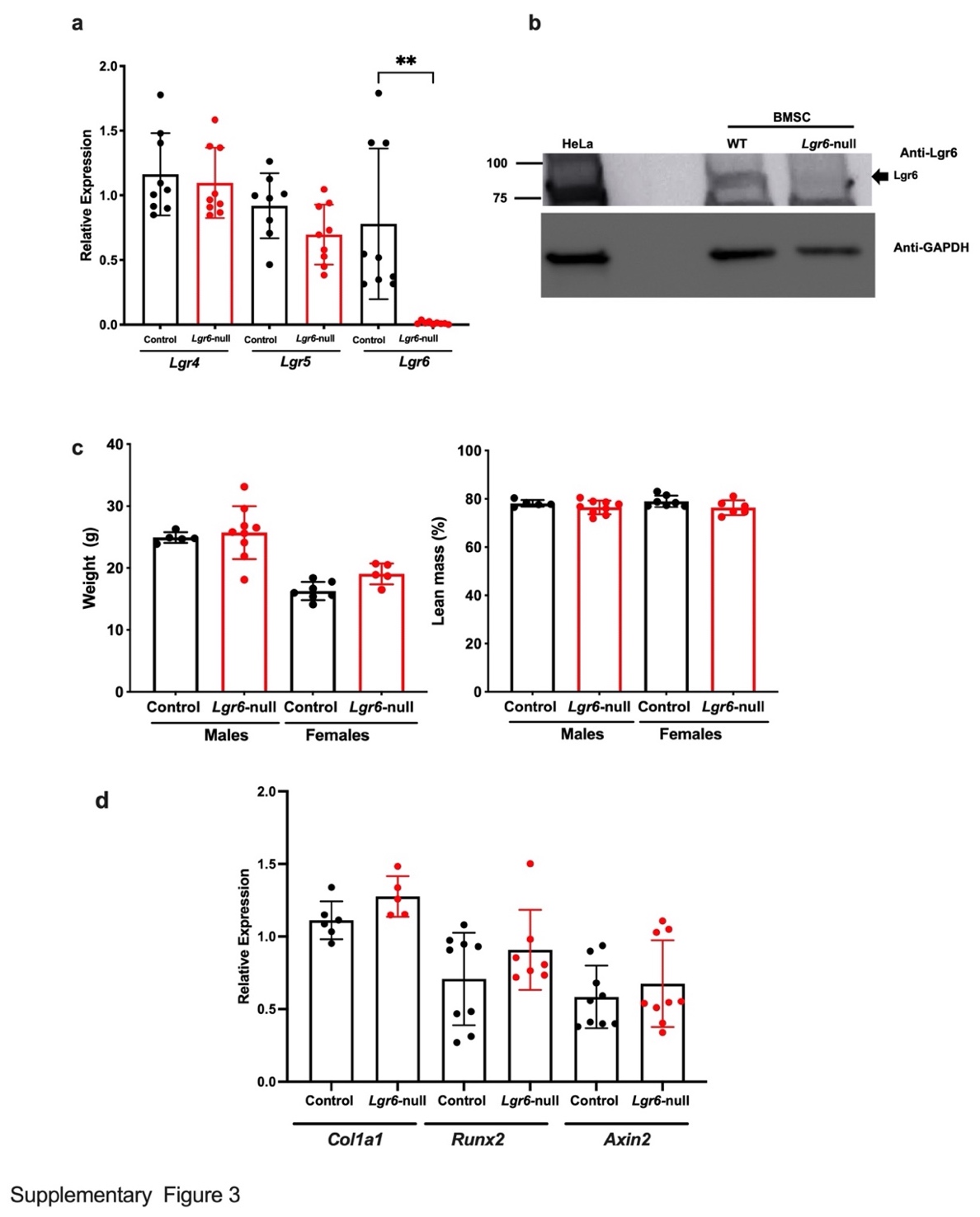

Supplementary Figure 1. Characterization of *Lgr6-*null mice. qRT-PCR analyses of Lgr family members. Expression of Lr6 protein was detected by Western blotting. HeLa cell lysate was used a positive control. No difference was observed in weight or lean mass of control of Lgr*6-*null mice. **c**. Expression levels of *Col1a1, Runx2, and Axin in* tibia were comparable between control and *Lgr6-*null.

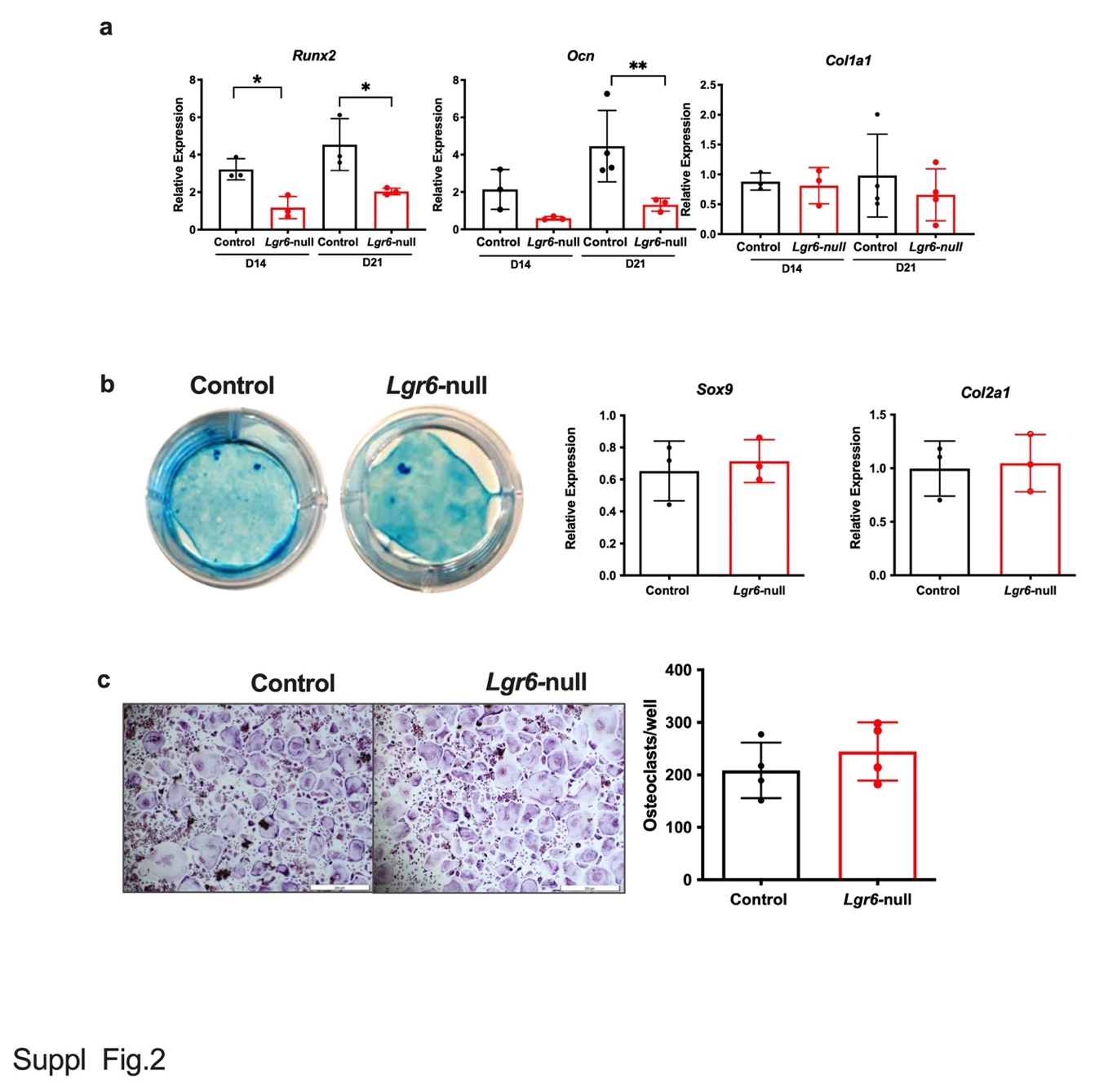

Supplementary Figure 2a: qRT-PCR analyses of critical osteogenic genes on days 14 and 21 of osteogenic differentiation of periosteal cultures. A representative of 4 experiments is shown **b.** Periosteal cells underwent chondrogenic differentiation in vitro and were stained with Alcian blue or subjected to qRT-PCR analyses. **c.** *In vitro* osteoclastogenesis assay of control and *Lgr6*-null-derived cells, stained for TRAP. TRAP+ cells with >3 nuclei were considered osteoclasts, and osteoclasts per well were counted manually. A representative of 2 experiments is shown
